# Mechanics and fate stochasticity shape stem cell distribution in tissues

**DOI:** 10.64898/2026.06.10.731353

**Authors:** Johannes C. Krämer, Edouard Hannezo, Jens Elgeti

## Abstract

Balancing cellular loss in tissues requires fine balance of cell proliferation and differentiation. In differentiated tissues consisting of a single cell type, a mechanical regulation of proliferation has been proposed to underlie growth-control and homeostatic steady-states. Yet, how tissues containing different cell types with distinct proliferation rates, mechanical interactions, and spatial self-organization retain robust homeostasis of cell proportions remains poorly understood. Here, we combine particle-based mechanical models of proliferative tissues with a classical hierarchy of stem, progenitor, and differentiated cells, undergoing stochastic fate choices, and show that mechanical feedback alone is sufficient to stabilize populations. We derive analytically and computationally a phase diagram of possible stable states, in particular those maintained either via slow and rare stem cells with short-lived progenitors or no stem cells and long-lived progenitors. Our simulations uncover that mechanical control of growth is sufficient, in the absence of any codes of adhesion or extrinsic niche signals, to cause stable spatial structures, with small stem cell clusters forming and maintaining dynamical renewal units. Our results demonstrate how complex spatial structures can emerge in minimal stochastic and mechanical simulations with impact to understand the homeostasis of multi-cellular systems.

## 1 Introduction

A hallmark of higher multicellular life is to control cell fate decisions, i. e. proliferation and differentiation, to balance cell loss and maintain a functional tissue composition and structure. This means that feedback mechanisms must exist, not only to ensure that cell proliferation exceeds cell loss during development and growth, but also that the two are exactly balanced at homeostasis. However, the mechanisms ensuring this balance remain a fundamental open question of cell biology and biophysics.

In recent years, mechanical interactions between cells have gotten increasing attention [1]. Because cell growth and division exerts compressive forces within a tissue, and because different cellular events like division or apoptosis have been shown to be mechano-sensitive, i.e. dependent on the mechanical state of the tissue [2, 3, 4, 5, 6, 7, 8, 9, 10, 11, 12, 13, 14, 15], mechanical feedback has emerged as a natural candidate mechanism to ensure homeostasis. Furthermore, these types of modelling approaches have predicted a number of non-trivial features, such as the visco-elastic properties of tissues in the presence of division and apoptosis [2, 16, 3, 17, 8], the competition between different tissues, interface dynamics, and the evolution of coexisting species [18, 19, 20, 21, 22].

However, these modelling approaches have so far largely concentrated on tissues made of a single cell type, or of different cell types that are independent from each others, i.e. no fate interconversion. This is a critical limitation, as homeostasis does not only entail the stability of cell numbers and density, but also the stable and correct proportions of different cell types. In particular, most adult tissues are maintained by adult stem cells [23, 24], which can both self-renew, and give rise to tissue specific, specialized cell types [25]. Despite the increasing insights in quantifying the dynamics of stem fate choices, for instance via lineage tracing methods or proliferation kinetic measurements, we still know comparatively little into the mechanisms of fate choices and the design principles leading to spatio-temporal coordination between cell types [26], beyond generic theoretical features common to large classes of models [27]. A first class involves include either cell-level asymmetric divisions, in which case the question of fate proportion stability becomes trivial, as it is hard-coded in the division patterns of each individual cells. However, how this would act in fluctuating environments exposed to external perturbations remains unclear, as the model lacks adaptability. Population-level asymmetry, characterized by a mix of symmetric and asymmetric stem cell fate choices [28], presents an alternative way to control cell fate, while being adaptable. In that case, fate outcomes must be balanced at the population level, yet, how this is mechanistically encoded remains poorly understood. Experimentally, a variety of biochemical pathways, like *Wnt, TP53*, or *RAS*, have been shown to play a crucial role in controlling the cell cycle, i.e. growth, division, but also differentiation, and deregulation of these pathways is found in carcinoma [29, 30, 31], but how to relate these to collective fate decisions remain unclear.

Here, we aim to generalize models of tissue growth with mechanical feedback to complex tissues with stochastic fate choices. We consider a generic hierarchy of stochastically dividing stem (S) and progenitor/transit-amplifying (T) cells, which in principle have infinite self-renewing capacity, as well as non-proliferative differentiated (D) cells. We utilize compartment-based analytic theory and stochastic particle-based simulations to derive and understand a phase diagram of possible parameter combinations leading to stable tissues. Using the simulations, we further explore the structure and dynamics of stochastically differentiating tissues with mechanical feedback. Interestingly, these are fundamentally shaped by the stochasticity of fate choices, which lead to stable clustering of stem cells, in stark contrast to a situations of deterministic fate choices, which had led to sharp repulsion and separation of single stem cells within the tissue [32]. Overall, we demonstrate that niche-like stem cell clusters can arise in a self-organized manner, simply as a result of an interplay between tissue mechanics and cell fate stochasticity.

## 2 Mechanical control of fate proportions in stemcell derived tissues

We consider a minimal model of complex tissues, where S and T cells symmetrically proliferate with rates *d*_*S*_ and *d*_*T*_, respectively. In addition to this, both populations can stochastically differentiate, respectively into T and D cells at rates *c*_*S*_ and *c*_*T*_ . D cells are post-mitotic, i.e. non-growing, cells and lost with constant rate *k*_*a*_. In the absence of feedback, such scenario is structurally unstable, and such model only has a steady state with extreme parameter fine-tuning (e.g. *d*_*S*_ = *c*_*S*_), and still prone to stochastic drift, as the rates do not adapt to the number of cells. Coupling the growth rates to be modulated by local mechanical pressure, solves this issue as we will see below. A schematic of this model can be found in Fig. 1 (A). While our model only includes symmetric divisions, the effective outcome of a division can be asymmetric, i. e. one cell undergoes another division whereas the other differentiates [33].

**Figure 1:**
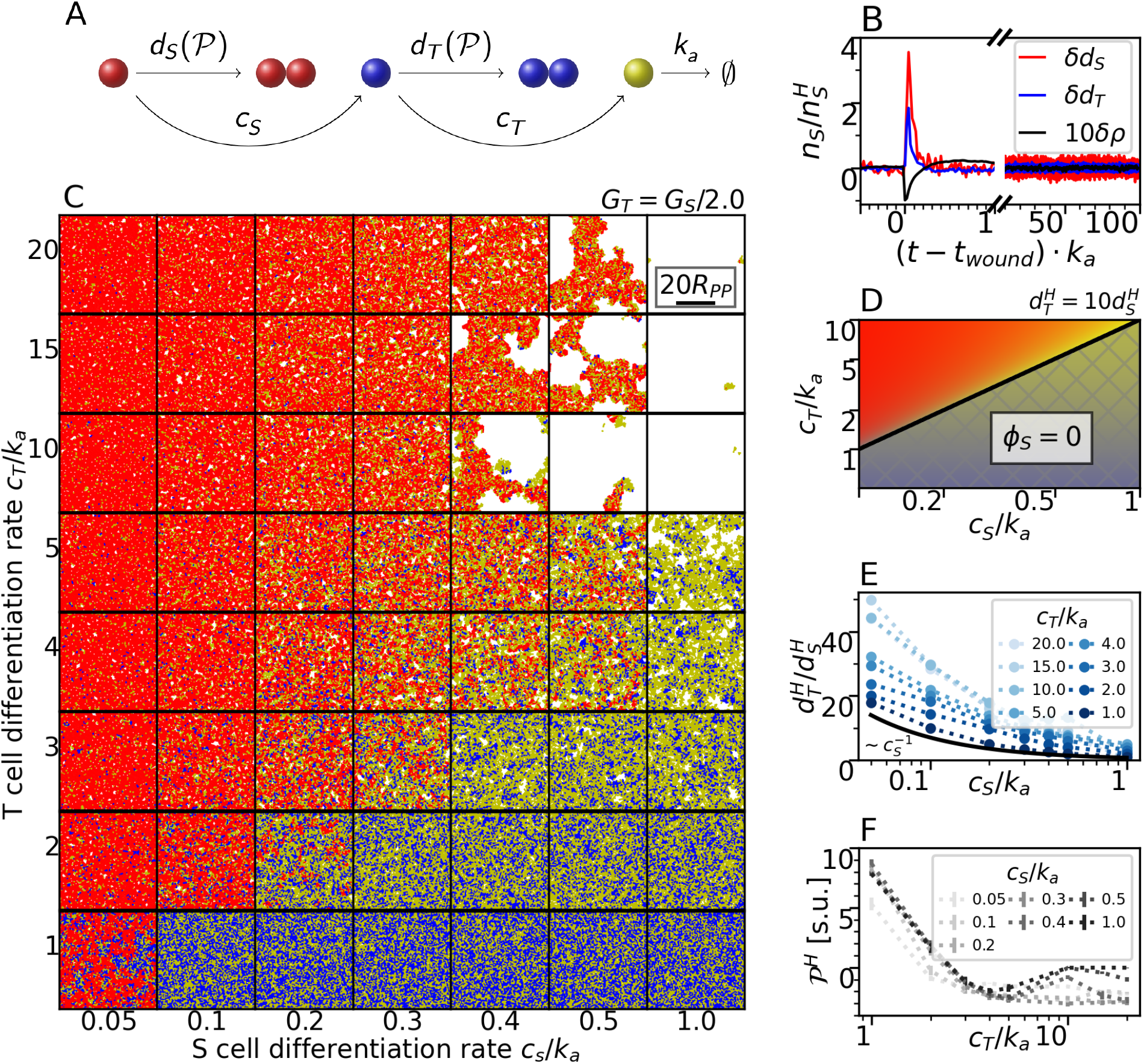
Homeostasis of S cell derived tissues via mechanical control. (A) Schematic representation of the S (red) T (blue) D (yellow) cell type model. This color code to specify cell type is applied throughout the whole work. (B) Response to a “wound”, i.e. at *t* = *t*_*wound*_, 10 % of cells are removed at random from the simulation of 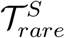 (def. in Fig. 2), where *δX* = *X*(*t*)*/X*^*H*^ − 1 for S and T cell division and total cell density (black, tenfold amplified for visibility) is shown. (C) Simulation snapshot phase diagram in steady state for variation of S and T cell differentiation rates *c*_(*S,T* )_ for fixed *G*_*T*_ */G*_*S*_ = 2. (D) Corresponding cell number fraction phase diagram following from Eqs. 2. Black line marks transition to S cell free region (hashed). (E) T cell division rate in homeostasis 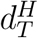 divided by 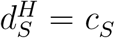 and (F) measured homeostatic pressure 𝒫^*H*^ in simulation units are shown for simulation phase space (B) as function of *c*_*S*_ and *c*_*T*_ .

We implement this lineage model into the two particle growth model, which considers cell mechanics and its feedback on divisions [34, 8, 32] (See Sec. 3). Thanks to the mechanical feedback on growth, we always find a stable steady state, independent of parameter tuning or initial conditions. Fig. 1 (B) shows that the mechanical feedback inherent to our simulations lies at the core of the stability: After randomly removing 10 % of the cells, division rates rise in response to lower cell density until the steady state is reached again (see also Animation S1).

The compositions and structure, however, varies greatly with cell type specific rates. Initially starting from a dense tissue with randomly assigned cell types, steady states with very different cell compositions, with S, T, or D cell rich regions (see Fig. 1 (C)) are found depending on the differentiation rates. This includes regions in which the S cell goes extinct but the tissue is continuously maintained by the transient cell type, as well as regions where the tissue loses integrity.

To better understand the different regions of the phase diagram and how to control the cell cycle dynamics to arrive at a biological relevant parameter range, we derive a coarse grained approximation, based on compartment models of each cell type [35], for cell fate proportions, starting with the dynamic equations describing the introduced model, given by

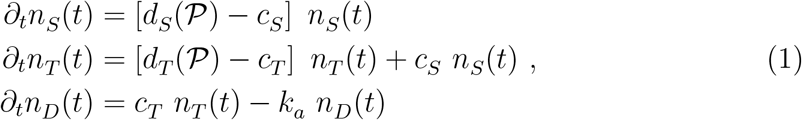

where *n*_*i*_(*t*) are the cell numbers of cell types *i* ∈ (*S, T, D*), and *d*_*S*_, *d*_*T*_, *c*_*S*_, *c*_*T*_, and *k*_*a*_ the division, differentiation, and loss rates, with explicit dependence on the system pressure 𝒫 in the division rates. Note that this dependence on pressure is just the lowest order expansion of the growth rate on mechanics [2]. We focus here on the steady state, i.e. the *homeostatic* state, where the numbers of cells of each type do not change over time (*∂*_*t*_*n*_*i*_ = 0). The steady state tissue pressure is referred to as *homeostatic pressure 𝒫*^*H*^ . Note that throughout this work the superscript *H* always defines the value of a variable in the homeostatic state.

From the dynamic equation for the S cell, we can then derive the homeostatic condition under which the S cell division rate balances its differentiation rate 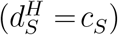, as a result of the mechanical feedback: Excess divisions will lead to an increase in pressure, which in turn will result in a decrease of divisions until numbers are back to their steady state value and the condition *d*_*S*_ = *c*_*S*_ is satisfied again. Steady state of T cells implies that the loss through differentiation is balanced by self-renewal plus influx of differentiating S cells. Similarly, the flux of D cells generated by T cell differentiation has to be balanced by loss of D cells. This allows us to derive the steady state cell number fractions 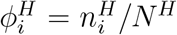, where 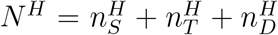, as functions of the differentiation and homeostatic division rates, given by

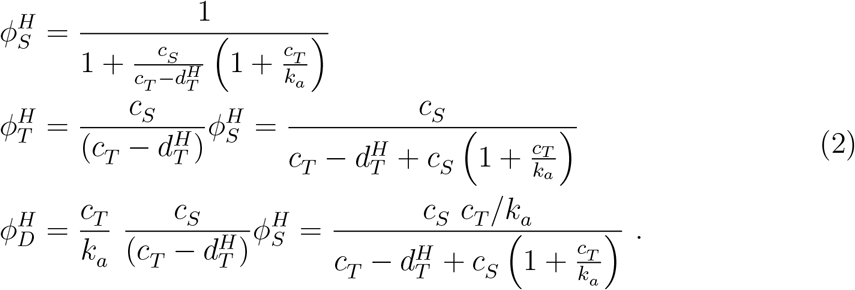

A phase diagram for cell number fractions as function of the S and T cell differentiation rate qualitatively displays the same trends as particle simulations as illustrated by simulation snapshots (see Fig. 1 B&D). Vice versa, we also derive the differentiation rates as functions of homeostatic variables (see SI Sec. S.3), which allows us to tune simulation parameters to match specific tissues. First, we envision a realistic tissue composition defined by set cell number fractions 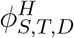 . Then, based on the measured or presumed division rates of S cells, which are typically slow cycling, and T cells, which divide frequently, Eq. 3 guide us with differentiation rates to use in the particle based simulations.

To go further, we also write linear expansions for the division rates *d*_*S*_ (𝒫), *d*_*T*_ (𝒫) as a function of pressure 𝒫, which describe how S and T cells adapt their division with respect to pressure variations and depend in general on the mechanical properties of cells and configuration of the tissue (see SI Sec. S.3). The homeostatic pressure is set when either 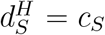 and 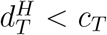 or 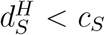 and 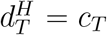. In the latter case S cells go extinct. In the case where S cells are present, the homeostatic pressure decreases with increasing S cell differentiation rate as also observed in our simulations (see Fig. 1 (F)). The homeostatic pressure can be negative for tensile tissues, but this has a lower bound. Below this limit, the tissue begins to rip and is losing confluency, as seen in the top right corner of Fig. 1 (C). For such non-confluent tissues, the global system pressure vanishes (𝒫= 0, see Fig.1 (F)). Similar analysis of the T cell division rates (see SI S.3) reveals more in depth insight in its effect on the cell composition, and we also demonstrate (see SI Fig. S1), how division rates can be controlled by changing the mechanical properties of cells, exemplarily shown by varying the T cell growth coefficient *G*_*T*_ .

## 3 Agent-based simulations of stem cell derived tissues

To go beyond understanding the average fate proportions of tissues, we then turned to particle-based simulations allowing us to understand stem cell dynamics in space and time. Following our previous work [32] we extend the two-particle growth model [34] to include stochastic cell fate. Here, we present only the essential aspects of the model for this work (see SI S.4 for full details). In essence, cells are modeled as soft attractive particles with stress-dependent growth. Differentiation and cell loss are modelled as instantaneous stochastic processes with fixed rates. This allows us to set directly *c*_*S*_, *c*_*T*_, and *k*_*a*_ in simulations, where *k*_*a*_ depicts the time scale of our simulations, i. e. we measure time in D cell generations.

While differentiation rates are explicit simulation parameter, the division rates are not. Instead, they follow from the interactions between particles, their mechanical properties and the tissue composition [34, 8, 21, 22], via the same mechanical control mechanism discussed in the previous section. We confirmed in simulations that this lead, as expected from the analytics, to robust stabilization of fate properties.

For instance, in Fig. 1, we show a phase diagram in the *c*_*T*_ − *c*_*S*_-plane, fitting well to the predicted phase behavior from number balance (Eq. 2), and we present further dependencies between parameters and observables in the SI. Taken together, this shows that our simulations are a versatile tool for simulating the mechanodynamics of stem cell derived tissues.

The main advantage of simulations lies in their ability to provide insight in the spatio-temporal dynamics of tissues, while the mean field equations presented above allow us to specifically tune the cell cycle dynamics in simulations to obtain biological relevant tissue compositions and dynamics. For the remainder of this work, we define two tissues (see Fig. 2, and Animations S2a and S2b): (i) The *stem cell rare tissue* 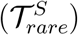 consists of approximately 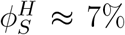 stem cells, 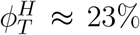 T cells and 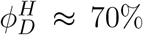 D cells. (ii) The *stem cell rich tissue* 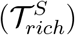 consists of approximately 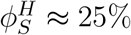 stem cells, 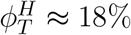 T cells and 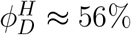 D cells.

**Figure 2:**
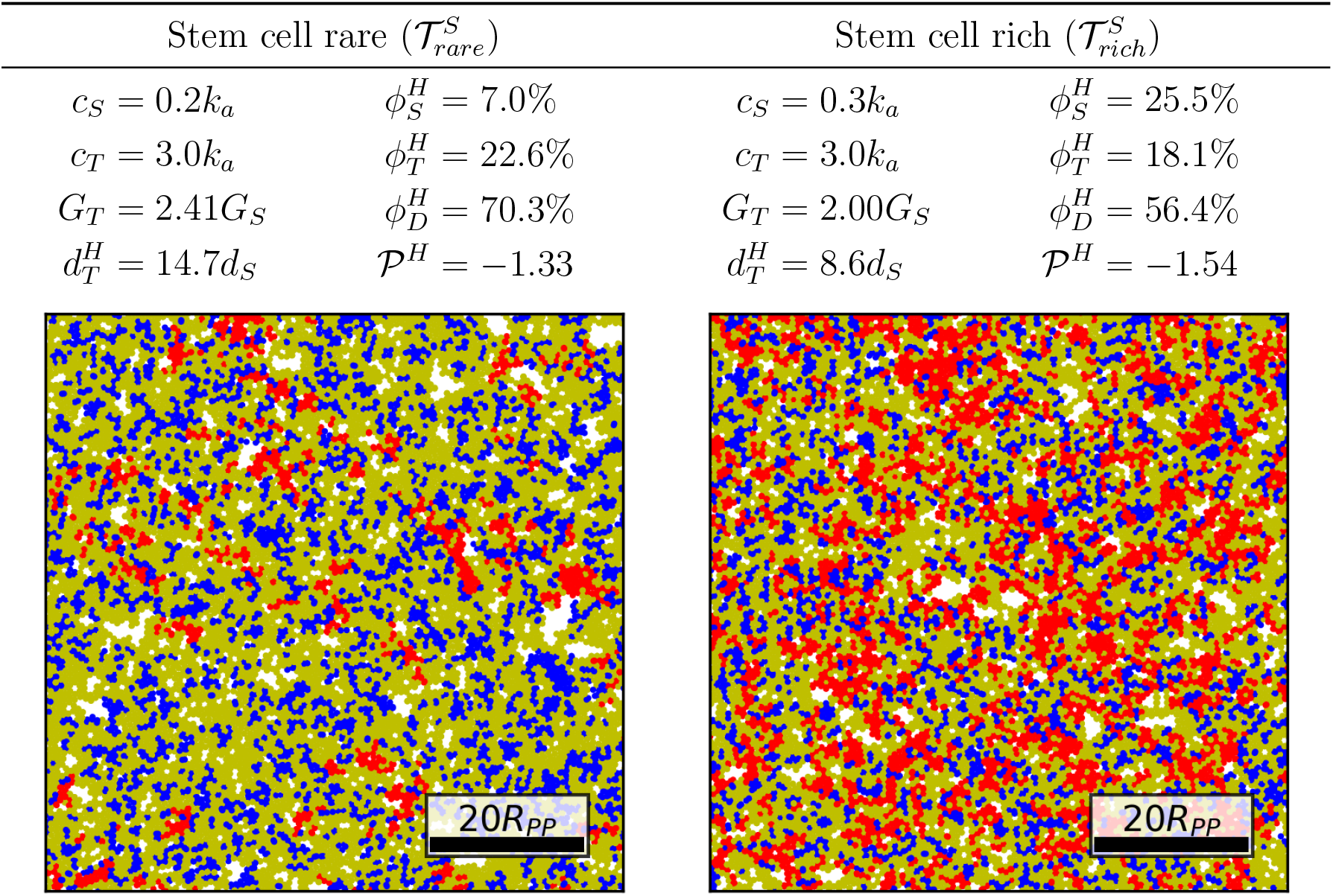
Details of selected tissues chosen for detailed analysis. Stem cell rare (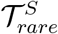, left) and rich (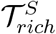, right) tissues studied in detail. Simulation parameters 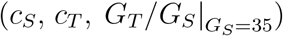 and resulting tissue properties 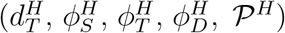 are given in the table together with a tissue snapshot in steady state. Stem (red), transient (blue), and differentiated (yellow) cells are drawn with cell size set to distance at which attractive and repulsive forces balance, and scale for 20 interaction radii *R*_*PP*_ is given.

In both tissues, stem cells are slow cycling: The measured T cell division rate is about one order of magnitude faster than the stem cells division rate 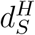 (see Fig. 1 (E)&2). These numbers are in the range reported for the interfollicular epidermis [36], in which stem cells divide 4-6 times per year and long-lived progenitor cells about once per week. Robustness of our results is confirmed by additional tissue configurations (see SI S.6), which cover a broader range of tissue compositions and cell fate rates but are not shown in the main text for the sake of clarity.

## 4 Self-organized formation of stem cell clusters in stem cell derived tissues

Interestingly, we find that that stem cells form well-defined spatial structures, with isolated niche-like stem cell clusters that remain relatively stationary in time, despite the fact that we have not implemented any specific adhesion between stem cells. Furthermore, this is in stark contrast both to previous works in the absence of mechanical interactions, where stem cell domains grow and phase separate over time [37], as well as our previous work with deterministic and asymmetrically dividing stem cells, which led to isolated stem cells due to effective and active proliferation-induced repulsion. This suggests that mechanical and stochastic effects cooperate to give rise to complex spatio-temporal patterns of finite-sized stem cell clusters, reminiscent of the concept of stem cell niches [38, 39], yet without any extrinsic spatially modulated cures from the substrate.

To systematically quantify our observed stem cell clusters, we use several metrics which could serve as guides for future quantification in experimental datasets. Firstly, we compute the stem cell pair correlation function *g*(*r*), which measures the relative density of S cells in the neighborhood of other S cells (see SI S.6.1). The niche like structure manifests itself in an increased S cell density in the vicinity of a S cell, i. e. within around ten cell sizes *R*_*PP*_ . For the low S cell fraction 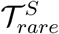, the integrated size of the peak 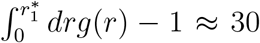, where the first root of 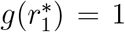 defines the upper bound, is a measure for the number of stem cells in a niche. The subsequent large minimum for small S cell fraction indicates an effective repulsion of S cell niches from the intermediate vicinity. The higher order peaks at even larger distances (*r* ∼ 50*R*_*PP*_ ) indicates a higher order structure formation of niches having well defined distances. This is further confirmed by a peak in the number fluctuations (see SI S.6). On the contrary, in the stem cell rich tissue 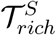, the pair correlation function decays much quicker, shows only a small minimum and is about unity thereafter, corresponding to limited S cell condensation with no significant further structure.

Next, we turned to the cluster size distribution. Defining all S cells within a distance of twice the cell diameter (2*R*_*PP*_ ) to belong to the same cluster, we find a mean cluster size of 17 cells for the S cell rare tissue, while for the S cell rich tissue, the mean cluster size reaches above 100, and we observe some huge, system spanning clusters.

Finally, we expand our observation to the study of the S cell-free regions. The void exclusion function *E*_*V*_ (*r*) measures the probability that at a random point has no S cell within a distance *r*. For randomly distributed particles, *E*_*V*_ = exp(−*ρπr*^2^). Indeed we find an exponential decay of *E*_*V*_ over a large range. However, the scaling is described by an effectively rescaled S cell density, due to clustering of S cells. For the S cell rare tissue, the prefactor corresponds to a much reduced S cell density of 0.8%, one order of magnitude smaller than the actual S cell density, consistent with the strong niche-like cluster formation discussed above. The percentiles of the void exclusion function across individual snapshots reveals further, that every snapshot contains at least one large void. Vice versa, the void exclusion function for the SC-rich tissue the exponential decay is less clear and roughly corresponds to densities of 6 − 14%, about half to one fourth of the actual S cell density. Here, the percentiles reveal that large voids are absent across the whole simulation.

Such local clustering of S cells and long range order is in stark contrast to asymmetrically dividing S cells with deterministic T cell duplication which results in an effective repulsion isolating individual stem cells, as we showed in our previous work [32]. Due to the symmetric divisions, an S cell pair is created next to each other, causing an increased probability to find two S cells at short distances before they separate from each other. The homeostatic control then causes long ranged self-organization of cells.

Finally, snapshots reveal another important feature: S cells are not dominantly surrounded by T cells as one might expect, and as we found in previous work with deterministic fate choices [32]. Indeed, quantifying (Fig. 3, see SI for method) the neighborhood composition reveals that T cells are even depleted from the S cell neighborhood. This is in stark comparisons to simulations that we performed with similar fate composition as 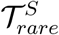 but with a deterministic differentiation scheme, where for slow and fast cycling stem cells, stem cells are largely surrounded by transient cells as expected. This underlines that inference of cell lineages and hierarchy from neighbourhood relations in static snapshots is prone to errors, that can be avoided by tracking dynamic relations.

**Figure 3:**
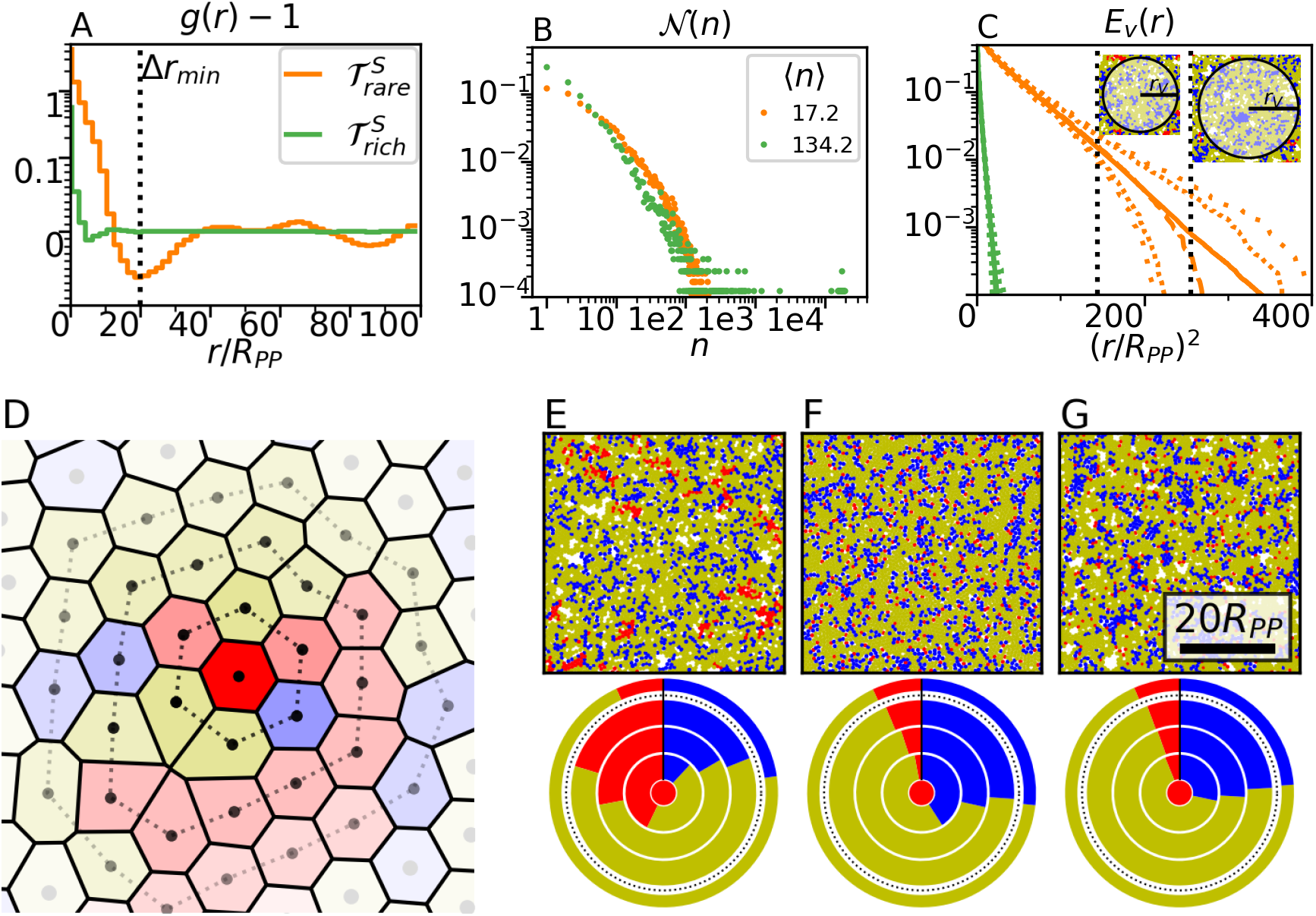
Spatial structure and niche formation. (A) Stem cell pair correlation function *g*(*r*) over pair distance *r* = |***r***_*i*_ − ***r***_*j*_|, with linear y-scale between (−0.1, 0.1) and logarithmic y-scale else. (B) Cluster size distribution function 𝒩 (*n*) for density based clustering with intra-cluster distance threshold 2*R*_*PP*_ . Average cluster sizes ⟨*n*⟩ are given in legend. (C) Void exclusion functions *E*_*V*_ (*r*) over squared distance (solid line for entire data set), following an exponential decay. Percentiles of *E*_*V*_ (*r*) of *individual* snapshots are shown in loosely dotted (3^*rd*^ *&* 97^*th*^), finely dotted (10^*th*^ & 90^*th*^), and dashed (50^*th*^) lines. Dashed vertical black lines denote example snapshots at void areas marked by dotted vertical lines. (D) Representative snapshot of the tissue used to construct cell neighborhoods. Cells of first, second, and third order are connected by dotted line and color opacity is gradually reduced with increasing order. (E-G) Snapshots (top row) of various differentiation models, (E) slow cycling S cells and stochastic differentiation 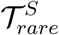, (F) fast cycling and (G) slow cycling stem cells with deterministic differentiation. For each system, corresponding neighborhood statistics (bottom row) are shown in pie charts around S cells (inner circle). Each ring represents neighbor statistics of first to third order and outermost ring, separated by dashed line, shows the global cell number fractions.

## 5 Long-term temporal dynamics of stem cell clusters

Beyond these static spatial correlations, simulations allow us direct access to several temporal dynamics of stem cell tissues, such as lineage histories or cluster stability over time. Simulation movies (see Animation S2a and S2b) and stem cell spatial distribution maps (see Fig. 4) suggest that S cell clusters form stable units, which fluctuate in size, yet only diffuse spatially at a very slow pace. The remarkable stability of S cell clusters for 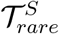 is highlighted by spatial distribution maps for S cell trajectories (see Fig. 4 A). Integrating stem cell densities, and displaying them over increasing time intervals shows, how clusters are remodeled but their overall structure remains unchanged for times much longer than the stem cell generation. This suggests that stable niche-like structures can be formed even in the presence of any external spatial signals.

**Figure 4:**
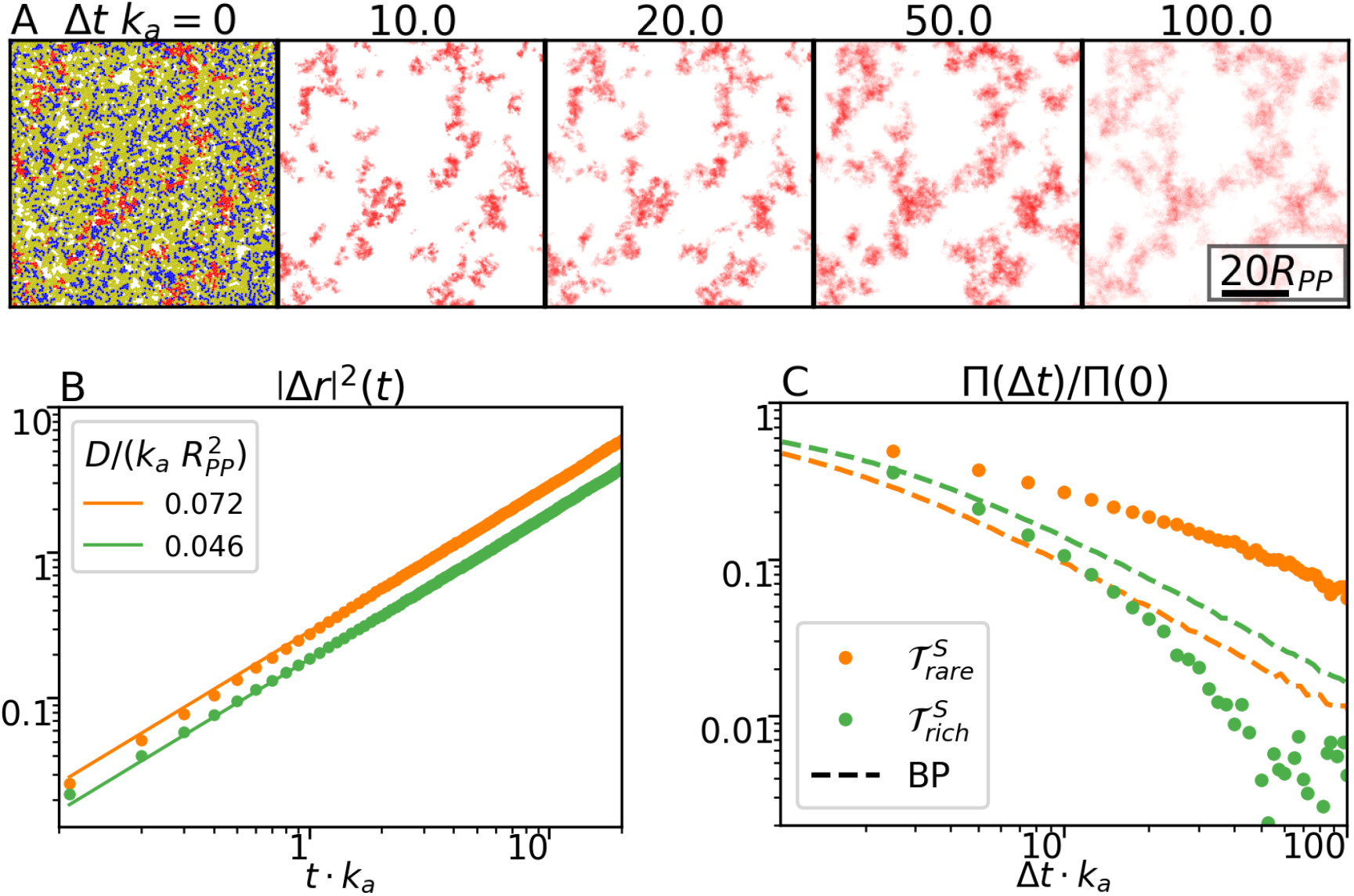
Temporal stability of stem-cell clusters. (A) Stem cell spatial distribution map following the initial configuration (left) at time lags of 10, 20, 50 and 100 D cell generations. Stem cell trajectories with opacity such that pixels, which are constantly occupied by an S cell, are fully red. Lighter or white regions correspond to low or no stem cell frequency, respectively. (B) Mean squared displacement of stem cells over time, measured from trajectories up to time of division or differentiation. (C) Density-fluctuation correlation function over time for tissue simulations (dots) and Brownian particle (BP) simulations (dashed line) with diffusion coefficient derived from (B) and BP number equal to mean stem cell number.

We quantify this remarkable stability by the autocorrelation function for density fluctuations of S cells, given by 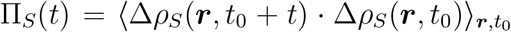, with 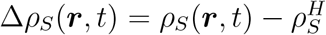 in subvolumes of size 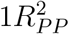 . As a guide, we compare the decay with a system of non-interacting Brownian particles, with diffusion coefficients derived from the single stem cell motion (see below). In the stem cell rare tissue, we find much slower decorrelation, evidencing more stable structures. On the other hand, in the stem cell rich tissue, density fluctuations decorrelate quicker than the passive system.

Additionally, we track individual S cells up to the time of division or differentiation, to quantify the mobility of individual cells. From this data, we calculate the SC mean squared displacement, which is found to be well described by passive diffusion |Δ*r*|^2^ = 2*dDt*, with dimension *d* = 2 (see Fig. 4). The diffusion coefficient corresponds to approximately half to one cell size per stem cell generation, consistent with previous simulations of homogeneous, mono-clonal tissues [16]. However, this is in stark contrast to our earlier work, in which we studied a hierarchical scheme with fixed number of symmetric T cell divisions and asymmetric S cell divisions only, where we found superdiffusion of the S cell induced by progeny propulsion. This points out, that the division-differentiation scheme does not only influence the structural properties, but also the dynamics of cells.

Finally, the ability to follow individual cell trajectories enables us to get detailed, time resolved information on the evolution of cell lineages. Similarly to lineage tracing experiments, we label cells in the homeostatic state at a specific time point, and trace their progenies over time. As expected from neutral drift, snapshots (see Fig. 5) reveal, that with time the total number of surviving clones decreases and the size of individual clones increases, as S cell clusters become monoclonal. Interestingly, S and T cells are non-trivially distributed, as clones can extend such that T cell contributions diverge from their S cell origin. This extends our previous, static, analysis and confirmed unexpected neighborhood distributions arising from stochasticity and mechanical interactions.

**Figure 5:**
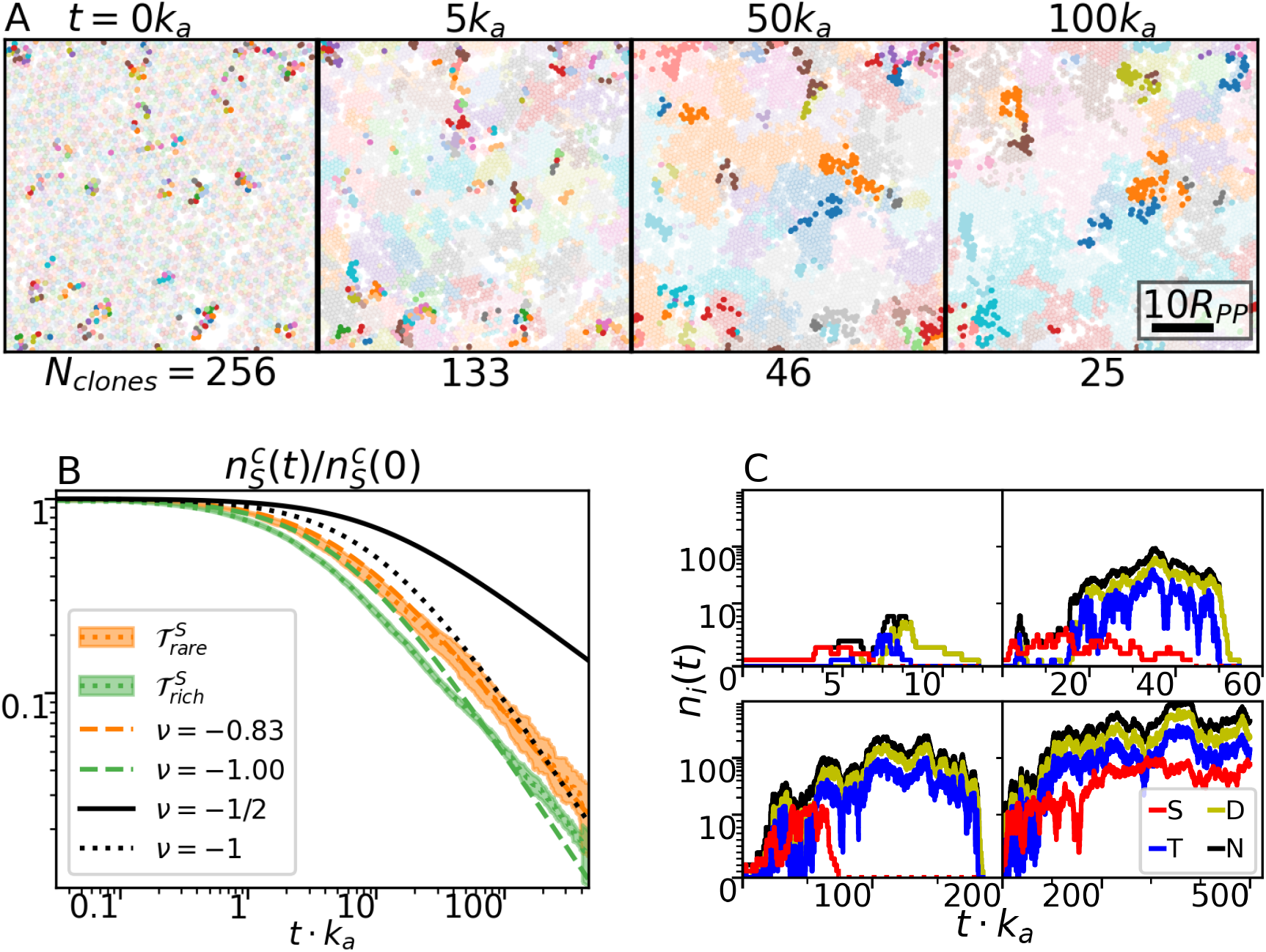
Lineage tracing in mechanical simulations. (A) Example of tissue clone evolution over time, where each clone is shown in different color, with reduced opacity for T and D cells to emphasize the S cell positions. Cells were marked with unique labels in the beginning and pass this over to daughter cells. (B) Fraction of surviving stem cell clones over time. Dotted line shows average over 12 independent runs and filled region standard deviation of clone fraction. (*c*_*S*_ · *t* + 1)^*ν*^ is shown for different exponents, given in legend, where dashed colored lines are best fit to data of 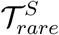 and 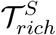. (C) Average number of clones per cluster over time for one short lived clone, two unlasting, and one surviving clone. Note the different time scales.

Measuring the clone survival fraction over time (Fig. 5B), we find that neutral drift theory, which predicts a decay of 1*/*(1+*c*_*s*_*t*)^*ν*^, with *ν* = 1 in two dimensions [40], does not fit the data well. Instead, we can describe the decay better with exponents *ν* = −0.63 and −0.76 for the stem cell rare and rich tissues, respectively, which hints to the fact that mechanical interactions create spatio-temporal structures that are not captured well by mean-field analytical approximations. We illustrate this by plotting the temporal evolution of specific clones in Fig. 5C. We find that while all clones which lose their stem cell eventually go extinct, they can exist for an extended period due to T cells, resulting in very long correlations not captured by simple neutral drift, a prediction which could be tested in experiments by combining intravital live-imaging with reporters of cell fate.

## 6 Discussion

Combining analytical theory and minimal mechanical simulations, we map a phase diagram of possible composition for cycling tissues composed of a hierarchy of stem cells, progenitors, and differentiated cells. We uncover how a mechanical regulation of growth can generically give rise to well-defined homeostatic states with different possible cell type fractions, and derive conditions for the stability of each cellular compartment. Agent based simulations confirm these theoretical predictions, but also provide novel insights in the spatial structure and temporal dynamics of stem cell self-organization. In particular, we find, that for low stem cell densities, stable and long range order emerges, even in the absence of external signaling. This order manifests as localized stem cell clusters which are well-spaced from each other and are very stable over time. Furthermore, stochasticity of fate choices, together with our mechanical regulation of growth, result in highly non-trivial neighborhood relationships, with progenitors being statistically further from stem cell clusters than differentiated cells, despite directly originating from stem cells. We contrast these findings with previous results not considering mechanical interactions [37, 41], as well as results without stochastic fate choices [32], which both show very different large-scale spatio-temporal dynamics. In this work, we remained agnostic about the molecular mechanisms underpinning how local pressure changes proliferation. However, wow biological cells sense and react to mechanics is an ongoing field of research, where recent findings suggest that membrane potential serves as a mechanotransductive sensor for pressure and thus regulates growth [15]. Further model extensions could therefore concentrate on understanding how specific models of mechano-sensing impact on global tissue dynamics.

Importantly, extensive experimental work has been done over recent years to extract cellular dynamics and fate choices from snapshots, for instance staining of putative fate markers or lineage tracing methods [26, 40]. For instance, the existence of stem cell clusters in epidermal cultures in vitro has been proposed to be linked to stem cell adhesion [42], although our simulations suggest they could arise without any specific adhesion code. Our neighborhood probability analysis and clone tracing (Fig. 3 (D-G)&5 (A)) also suggests that caution should be taken in simply assuming that neighboring cells are immediate progenies of stem cells, in particular in noisy systems [43]. Overall, combining our simulations with static experimental data should allow to infer, from detailed spatial correlation and tissue structure, some of the key microscopic details of the underlying division and differentiation schemes, which poses a valuable tool to test biological hypotheses. Finally, our simulations also make a number of detailed predictions on the dynamics of stem cell clusters and organization within cycling tissues. Over recent years, advances in intravital imaging have allowed unprecedented insights into the dynamics of renewal, for instance in epidermis or intestine [44, 45, 46, 47, 48, 49]. In particular when combined with live-reporters of cell fate or state, these datasets would be ideal to test our experimental findings, in particular temporal correlations between fate transition and local mechanical properties or relative cell densities, as well as the temporal stability of clusters of different cell types. Recent literature has also shown that different cell types within the same tissue are endowed with very different mechanical properties [10, 50, 51, 52], From a more theoretical perspective, potential future extension of our model could be to generalize it to 3D tissues, for instance skin where differentiation is coupled to stratification to upper layers [50], consider models with cell fate reversibility [37], which can be more flexibility to cellular responses.

Finally, considering in our simulations the role of interactions between cells and their micro environment, for instance substrate geometry or stiffness, would likely lead to interesting phase diagram which could be further compared with experiments [53, 54, 55, 56].

## S Supplementary information (SI)

### S.1. Data availability

All original code has been deposited at Zenodo at https://doi.org/10.5281/zenodo.18245707 and **will be/is** publicly available as of the date of publication. Temporary access link during review: https://zenodo.org/records/18245707?preview=1&token=eyJhbGciOiJIUzUxMiIsImlhdCI6MTc3OTA5NzU5MCwiZXhwIjoxNzg1MzY5NTk5fQ.eyJpZCI6ImE1YTQ1MjJlLWEyOTQtNDE4NC05NGQ1LTk0YmRjYjBlYzZiNyIsImRhdGEiOnt9LCJyYW5kb20iOiJiZDlkN2IwNDJiODRiMDQzZWU3MTE0ODJmZjNkMmUyYSJ9.IoPw23n_jsBpcyWCw-J2_J2q3b9u_c2RcDjmcMITYS7Nq9X0MhfUg7rAiBK-P7Gn64Jr4iaHpEbFIHsN289kTA

### S.2. Supplementary Animations

We provide demonstrative animations of selected simulations:

1. *S1-TissueAnimation-RandomCellLoss*: animated simulation snapshots of stem cell rich tissue 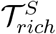 with 10% random cell loss.
2. *S2a-TissueAnimation-SCrare*: animated simulation snapshots of 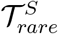
3. *S2a-TissueAnimation-SCrich*: animated simulation snapshots of 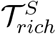

### S.3. Auxiliary information for continuum theory

#### Setting differentiation rates

In order to tune cell number fractions, we have derived the dependence of 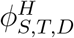 on cell fate rates. In simulations, however, we aim to tune the cell fate rates with a certain configuration in mind. Thus, we rewrite Eq. 2 to derive equations for the differentiation rates.

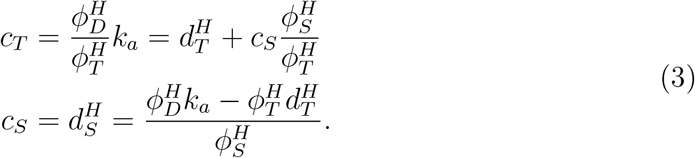

Note that the T cell division rate is not a simulation parameter, but follows the mechanical properties of cells, and thus requires accurate adjustment of parameters like for example the growth coefficients *G*_*S*.*T*_ .

#### Pressure dependence of division rates

In first order, we can expand the S and T cell division rates as a function of pressure 𝒫 as

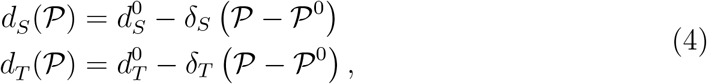

where 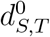 is the division rate of the system at pressure 𝒫^0^, an arbitrary expansion point. In homeostasis (*c*_*S*_ = *d*_*S*_), we consequently can rewrite the first equation as 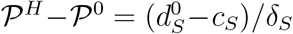 . Inserting this into *d*_*T*_ (𝒫^*H*^ ) yields 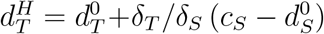 .

#### Average number of T cell cycles

In Sec. 4 and Fig. 3 D-G we compare the stochastic differentiation model with a deterministic differentiation model. In the latter, the number of T cell cycles is set by a hard-coded variable *N*_*T*_, such that T cells, which have divided *N*_*T*_ times, terminally differentiate automatically. In order to compare both systems, we need to chose an integer *N*_*T*_, that results in comparable cell number fractions. The number of T cells produced per S cell division in the stochastic model can be derived from the T cell number balance equation 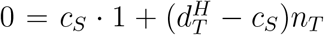 . Consequently, 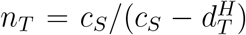 T cells are produced per S cell division. In the deterministic model, on the other hand, we have shown that each stem cell division can produce 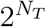 T cells. Thus, we have to set 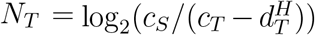, where *dT*^*H*^ is measured in simulations. For 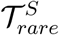, we set *N*_*T*_ = 6 in the deterministic model with slow cycling stem cells. For fast cycling stem cells, we cannot estimate the number, but have to test it via simulations, and find that *N*_*T*_ = 3 results in comparable cell number fractions.

### S.4 Two particle growth model (2PG) and simulation setup

The different stem cell differentiation models described in the main text are implemented in the two-particle growth (2PG) model of Refs. [34, 8] and has been adapted in Refs. [20, 21, 22, 32]. In the following, we requote the description already presented in [32] and extend it where needed.

Each cell is described by two particles, which repel each other via an active growth force

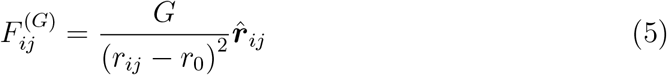

with unit vector 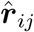, distance *r*_*ij*_ between the two particles, a constant *r*_0_, and growth strength *G*. For non-growing cells, like D cells, the growth strength is set to zero, *G*_*D*_ = 0.

To prevent overlap of cells, particles of different cells interact via a soft repulsive volume-exclusion force

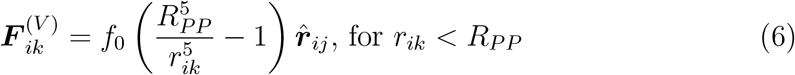

with exclusion strength *f*_0_ and interaction length *R*_*PP*_, which sets the length scale of our simulations. Further, cells in contact interact via an attractive adhesion force of the form

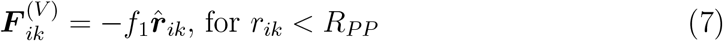

with adhesion strength *f*_1_.

We employ a dissipative particle dynamics-type thermostat [57], with an effective temperature *T*, to account for energy dissipation

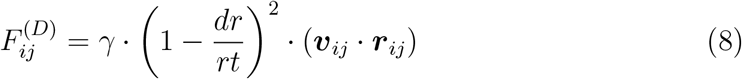

and random fluctuations

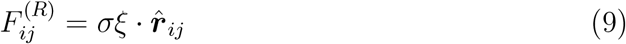

where *γ* and *σ* are related to fulfill the fluctuation-dissipation theorem [58]. Also, background dissipation is taken into account as

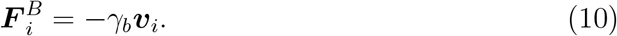

**Table S1:** Standard parameter set of agent based tissue simulations.

| Parameter | Symbol | Value |
| --- | --- | --- |
| Time step | $\Delta t$ | 0.001 |
| Pair potential interaction range | $R_{PP}$ | 1 |
| Cell expansion constant | $r_0$ | 1 |
| Division distance threshold | $R_{ct}$ | 0.8 |
| New cell particle initial distance | $R_d$ | $10^{-5}$ |
| Mass | $m$ | 1 |
| Intra-cell dissipation coefficient | $\gamma_c^{ST}, \gamma_c^D$ | $2G_{ST}, 2G_T$ |
| Inter-cell dissipation coefficient | $\gamma_t$ | 50 |
| Background dissipation coefficient | $\gamma_c$ | 0.1 |
| Apoptosis rate | $k_a$ | 0.01 |
| Noise intensity | $k_B T$ | 0.1 |
| Repulsive cell-cell potential coefficient | $f_0$ | 2.39566 |
| Attractive cell-cell potential coefficient | $f_1$ | 10.0 |

A self-consistent velocity-Verlet algorithm [59] is implemented to integrate the equations of motion, and all simulations were performed with periodic boundary conditions in 2D.

Cell division is performed, when the two particles of one cell are separated by a critical threshold *R*_*ct*_. A new particle is randomly placed near each original particle in a distance *R*_*d*_, and the two new particle pairs form the two daughter cells. For the model described in Fig. 1, differentiation is implemented as stochastic process with fixed rates *c*_*S*_ and *c*_*T*_ . In the deterministic model discussed in Sec. 4, differentiation takes place at the time of division, where for T cells an internal variable tracks the number of divisions and triggers terminal differentiation, when the maximum number of divisions is reached.

D cells are stochastically removed from the simulation with a finite apoptosis rate *k*_*a*_. Time is measured in terms of the inverse apoptosis rate, and referred to as “generation”.

In addition to the tissue specific parameters given in the table in Fig. 2, the standard parameter set for our simulations are given in Tab. S1. However, not all of these simulation parameters, have a direct conversion to physical units. As discussed in [34] one has to chose well defined measurable quantities, such as apoptosis rate and range of the pair potential as rescaling units for inverse time and particle diameter, to allow for conversion to physical units and comparison with experiments.

As simulation for large areas have high computational cost, we performed the screening of the parameter space in smaller areas of size 50*R*_*PP*_ × 50*R*_*PP*_ for 500 generations, starting with a random cell configuration with 10 % stem cells. Large tissues of size 200*R*_*PP*_ × 200*R*_*PP*_ were simulated by patches of these smaller simulations. Care has been taken, that patches are not correlated, when stitched together, and that the system equilibrates before measurements are conducted (see below).

#### S.4.1 Brownian dynamics simulations

In the main text (Sec. 5), we compare results obtained with the agent based model with a passive Brownian particle (BP) model to understand the difference in static and dynamic properties. The BP model diffusion is simulated by solving the differential equation

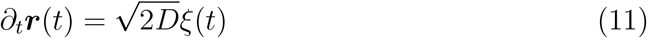

with diffusion coefficient *D* obtained from measurements of the stem cell MSD in the 2PG model and *ξ* describing white noise, which is mean centered ⟨*ξ*(*t*)⟩_*t*_ = 0 and uncorrelated ⟨*ξ*(*t*)*ξ*(*t*^*′*^)⟩ = *δ*(*t* − *t*^*′*^).

### S.5. Parameter selection: Growth rate variation

**Figure S1:**
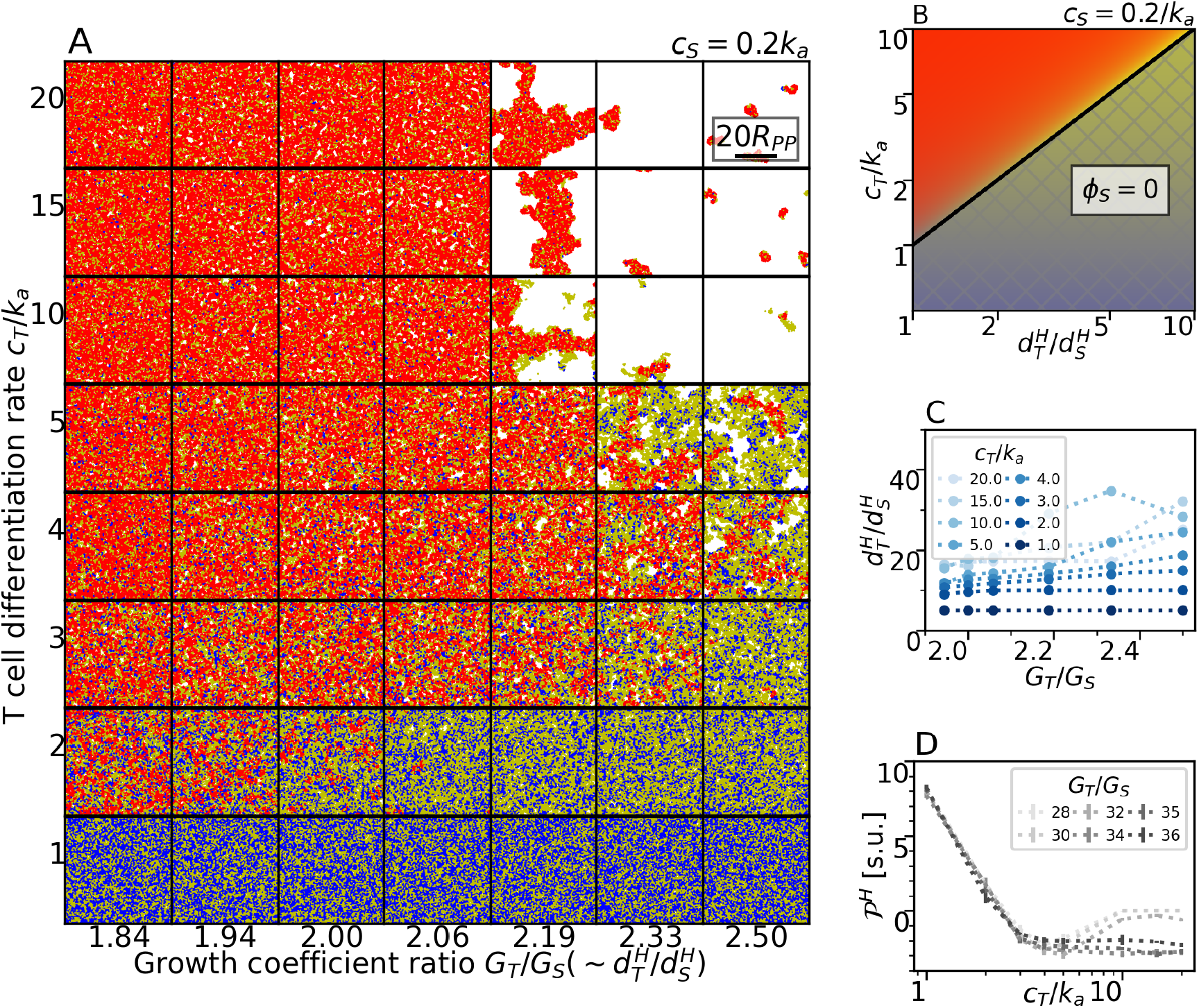
(A) Simulation snapshot phase space for *G*_*S*_ − *c*_*T*_ variation, (B) corresponding phase diagram for cell number fractions from Eq. 2, (C) T cell division rate, and (D) homeostatic pressure for measurements in (A).

In Sec. 2 of the main text, we showed how the differentiation rates of S and T cells change the cell number fractions. Further, the theory predicts that also the T cell division rate *d*_*T*_ does affect these ratios. Above, we have seen, that mechano-response (*δ*_*S,T*_ ) of different cell types shows mutual interference. To highlight that division rates adjust as a consequence of the mechanical interactions, we vary the stem cell growth coefficient, keeping *G*_*T*_ and 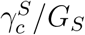 constant. Again, we find good agreement of simulations and prediction from the cell number balance equations (see Fig. S1 (A)&(B)), and how *G*_*S*_ allows to fine tune the T cell division rate, due to change in pressure. Also, division rates and pressure change in the same fashion as discussed in the main text (see Fig. S1 (C)&(D)).

**Figure S2:**
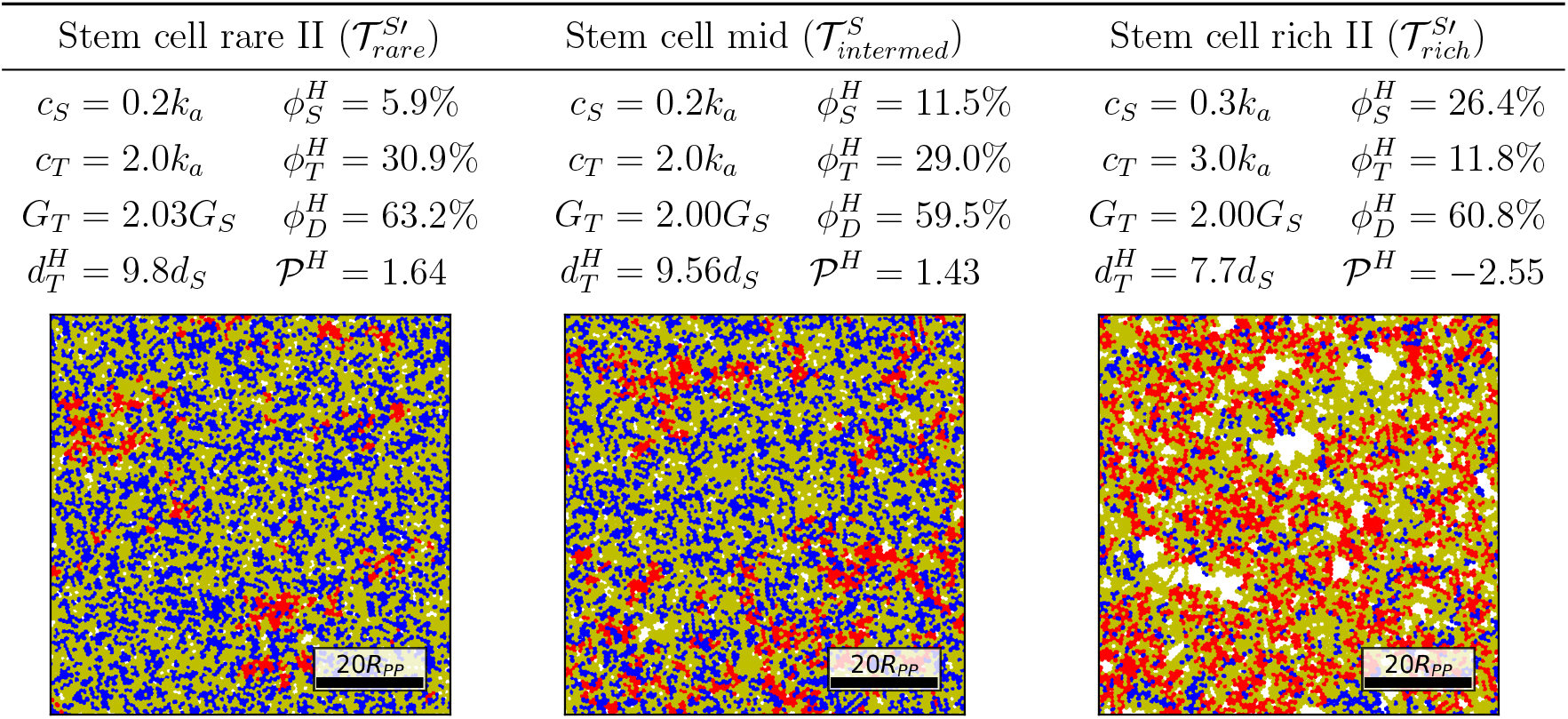
Additional stem cell rare (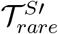, left) and rich (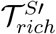, right) tissues and an intermediate tissue (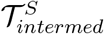, middle) to show robustness of results. Simulation parameters (*c*_*S*_, *c*_*T*_, *G*_*T*_ */G*_*S*_) and resulting tissue properties 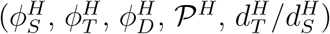 are given in the table together with a tissue snapshot in steady state. Stem (red), transient (blue), and differentiated (yellow) cells are drawn with cell size set to distance at which attractive and repulsive forces balance, and scale for 20 interaction radii *R*_*PP*_ is given.

### S.6. Additional tissue configurations to show robustness of main text results and observables

We utilize several observables, which we define here in addition with results for three more tissues, to show the robustness of our results (see Fig. S2). Beside another stem cell rare 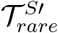 and rich 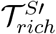 tissue, we also select a tissue of intermediate stem cell fraction. The intermediate tissue is very similar to 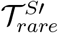, only differing only in the stem cell growth coefficient by less than 2%, which already leads to a doubling of the S cell fraction.

#### S.6.1 Static properties

##### Pair correlation function

*g*(*r*) The pair correlation function measures the likelihood to find two particles at a distance *r*. It is defined as

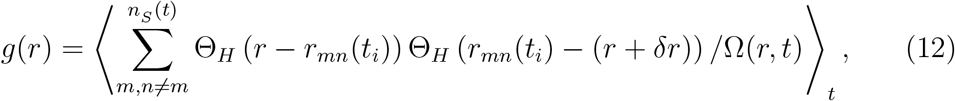

where *r*_*mn*_(*t*_*i*_) = |***r***_*m*_(*t*_*i*_) − ***r***_*n*_(*t*_*i*_)| is the pair distance, using the center of mass of cells, Θ_*H*_ Heaviside functions, used to construct a measurement annulus of width *δr*, and Ω a normalization factor given by

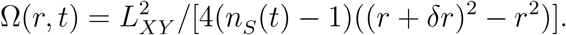

For randomly distributed particles *g*(*r*) = 1, which is why we display *g*(*r*) − 1 in Fig. 2 (A)+S3 (A). While we employ the pair correlation function for stem cell pairs, it can be generally be used to measure the pair correlation of other particle types, or cross type correlations.

**Figure S3:**
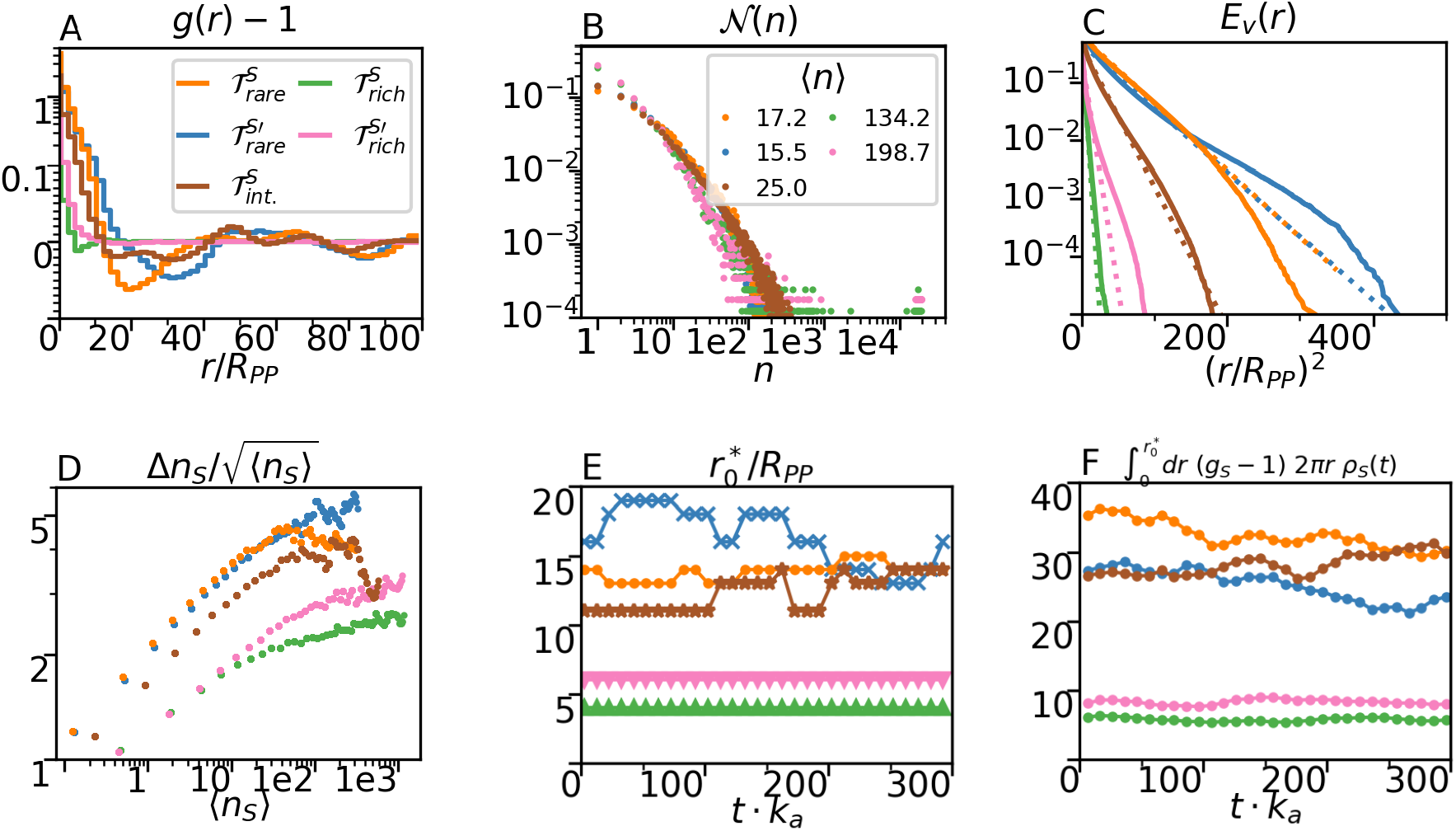
Niche formation in tissues of different stem cell densities, as described in the legend in subplot (A). (A) Pair correlation function, (B) cluster size distribution function, and (C) void exclusion function as shown in Fig. S3 in the main text. (D) shows the number fluctuations over mean cell numbers and (E) the first root of *g*(*r*) − 1 and (F) the integral of (*g*(*r*) − 1)2*πrρ*(*r*) from zero to this value.

In both stem cell rare tissues, and also in the tissue with intermediate stem cell fraction, we find a strong accumulation at short distances and depletion afterwards. However, the process is not as distinct for intermediate stem cell fraction, suggesting that niche formation is a feature which gets more pronounced with reduced stem cell fraction.

To quantify the short range excess of particles, we calculate the integral

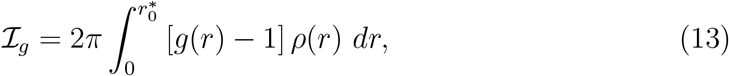

where 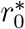 is the first root of *g*(*r*) = 1 and *ρ*(*r*) the particle density, i.e. stem cell density.

Both, the first root 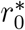 and integral ℐ_*g*_ are shown as function of time, i.e. calculated for individual snapshots, in Fig. S3 (E)&(F). This acts as measure of coarsening of the system, i.e. how structure shapes over time. We do not find any variation for the stem cell rich tissues. In stem cell rare tissues, we do see small changes on very long times caused by the steady state fluctuations, which become more visible at reduced S cell densities. However, there is no clear trend towards a coarsening of structures, as independent simulations confirm. Also, the intermediate configuration does not show clear indications for coarsening on the already long times simulated for all tissues. While the presented data to confirm absence of coarsening is shown from aggregated S cell distance data of four subsequent frames, spanning 10 D cell generations, we also looked in data of each independent snapshot and could not find any difference, except that noise increases for lower statistics.

##### Clustering

We calculate the cluster size distribution function

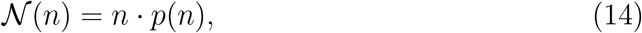

where *n* is the cluster size and *p*(*n*) the probability to find a cluster of size *n* (see Fig. S3 (B)). Clusters are searched using the `DBSCAN` algorithm with a cluster distance threshold of 2*R*_*PP*_ as implemented in [60].

Similar to the effective repulsion quantified via the pair correlation function, cluster sizes do increase with stem cell fraction, until percolated structures can be found for the stem cell rich tissues. For higher stem cell fractions, percolation eventually leads to the absence of isolated stem cells. However, clusters sizes are not proportional to the stem cell fraction and grow slower in size.

##### Void exclusion function

All previously discussed quantities are measured from particle positions. In simulations, however, we find that in particular the existence of large stem cell free regions is a remarkable feature. To quantify this, we measure the distance to the closest stem cell from a randomly chosen point. For each snapshot, we chose as many random points as stem cells and from the distance array construct the complementary cumulative distribution function (*cCDF*_*V*_, see Fig. S3 (C)). In Fig. 2, we show the percentiles for individual snapshots, and in addition calculate the *cCDF*_*V*_ from the distances of all snapshots combined to highlight how the analysis of individual snapshots can differ at low S cell densities.

##### Number fluctuations

Accumulation of SCs can further be analyzed by the number fluctuations: For increasing size of square subvolumes, we measure the particle number and its variance. Then, the variance divided by the square root of the mean number is plotted over the mean particle number (see Fig. S3 (D)). When particles are randomly distributed, the variance is expected to grow with the square root of the mean number, and thus should be constant at unity. In tissue simulations, we do find that stem cell number fluctuations are increased by a factor up to approximately five. Above subvolume lengths of approximately 20*R*_*PP*_, fluctuations begin to flatten. The broader and lower peak for higher SC-densities in turn is consistent with limited aggregation of SCs. Generally, the increase of fluctuations on short lengths and plateau at larger distances supports our finding of long range order and global homeostasis.

##### Neighborhood analysis

We show simulation results for different microscopic models, which all result in the same homeostatic cell number fractions (see Sec.4 - Fig. 2). However, the local configurations are not the same. This is quantified via a neighborhood likelihood composition. First, we perform a Delaunay triangulation, i.e. a graph connecting cell centers, for all cells to construct a neighbor list for each cell. Then, we recursively generate nearest, next-nearest, and second-next-nearest neighbor lists. To avoid overcounting for higher order neighbors, we utilize an exclusion list, in which we store the cells already counted at lower order. Finally, we merge these neighbor lists to calculate the average fraction of S, T, and D cells in first to third order around stem cells. Data from all stem cells of multiple simulation frames is aggregated. To visualize the results, we select an exemplary tissue section and visualize cells via a Voronoi diagram with the usual color coding and increasing shading for higher order neighbors.

**Figure S4:**
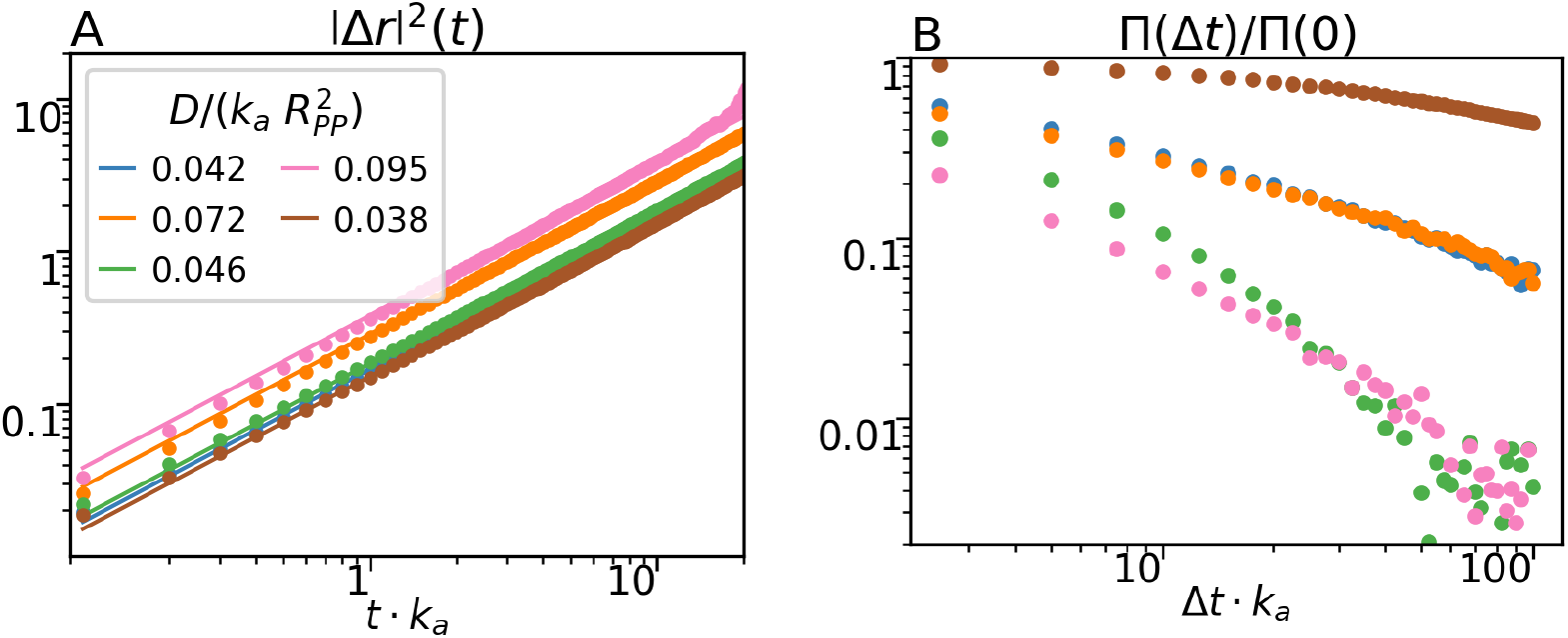
(A) Stem cell dynamics (MSD) and (B) stem cell density fluctuation correlation function.

#### S.6.2. Dynamic properties

##### Mean squared displacement

For each tissue configuration, we find the mean squared displacement in the same order of magnitude, around half a cell diameter per S cell generation (see Fig. S4 (A)). Also, in each tissue a linear slope describes the data well, confirming that no superdiffusive behavior is found.

##### Density fluctuation autocorrelation and conditional probability

From the stem cell trajectories, we measure local stem cell densities in small boxes of edge length 1*R*_*PP*_ . Then, we calculate the difference 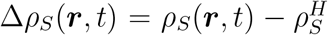 between local *ρ*_*S*_(***r***, *t*) and global 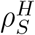 stem cell density for all subvolumes of the whole simulation volume (see Fig. S4 (B)).

We find the most stable stem cell niches for the intermediate tissue. In contrast to the stem cell rare tissues, the slight increase in stem cell density paired with slow dynamics seems to stabilize niches. Stem cell rich tissues show fast decorrelation, but also have the fastest differentiation dynamics.

